# Molecular properties and evolutionary origins of a parvovirus-derived myosin fusion gene in guinea pigs

**DOI:** 10.1101/572735

**Authors:** Ignacio Valencia-Herrera, Eduardo Cena-Ahumada, Fernando Faunes, Rodrigo Ibarra-Karmy, Robert J. Gifford, Gloria Arriagada

## Abstract

Sequences derived from parvoviruses (family *Parvoviridae*) are relatively common in animal genomes, but the functional significance of these endogenous parvoviral element (EPV) sequences remains unclear. In this study we use a combination of *in silico* and molecular biological approaches to investigate a fusion gene encoded by guinea pigs (genus *Cavia*) that is partially derived from an EPV. This gene, named *enRep-Myo9*, encodes a predicted polypeptide gene product comprising a partial *myosin*9 (*Myo9*)-like gene fused to a 3’ truncated, EPV- encoded replicase. We first examined the genomic and phylogenetic characteristics of the EPV locus that encodes the viral portions of *enRep-Myo9*. We show that this locus, named enRep, is specific to guinea pigs and derives from an ancient representative of the parvoviral genus *Dependoparvovirus* that integrated into the guinea pig germline 22-35 million years ago. Despite these ancient origins, however, the regions of enRep that are incorporated into the coding sequence of the *enRep-Myo9* gene are conserved across multiple species in the family Caviidae (guinea pigs and cavies) consistent with purifying selection. Using molecular biological approaches, we further demonstrate that: (i) *enRep-Myo9* mRNA is broadly transcribed in guinea pig cells; (ii) the cloned *enRep-Myo9* transcript can express a protein of the expected size in guinea pig cells *in vitro*, and; (iii) the expressed protein localizes to the cytosol. Our findings demonstrate that, consistent with a functional role, the *enRep-Myo9* fusion gene is evolutionarily conserved, broadly transcribed, and capable of expressing protein.

**Importance:** DNA from viruses has been ‘horizontally transferred’ to mammalian genomes during evolution, but the impact of this process on mammalian biology remains poorly understood. The findings of our study indicate that in guinea pigs a novel gene has evolved through fusion of host and virus genes.

## Introduction

Animal genomes contain numerous DNA sequences derived from viruses. These endogenous viral elements (EVEs) are thought to arise when infection of germline cells (i.e. gametes or early embryonic cells) lead to viral sequences becoming integrated into the chromosomal DNA of ancestral organism, such that they were subsequently inherited from parent to offspring as novel genes [1]. Examining EVE sequences has provided unexpected insights into the evolution of viruses and their hosts. EVEs preserve information about extremely ancient viruses, and can provide unique insight into the long-term evolutionary interactions between viruses and cells [2]. Furthermore, genomic and experimental research has revealed that during eukaryotic evolution, some of these ‘horizontally transferred’ viral sequences have been co-opted or ‘exapted’ by host species genomes, such they now perform physiologically relevant functions - for example, mammalian genomes contain EVEs with roles in cell function, embryonic development, and antiviral immunity [3-7].

Most of the EVE sequences found in mammalian genomes are derived from retroviruses (family *Retroviridae*). However, EVEs derived from several non-retroviral virus families have also been identified, and among these, parvoviruses are well represented [8-10]. Parvoviruses (family *Parvoviridae*) are small, non-enveloped viruses of icosahedral symmetry. They have a linear, single-stranded DNA (ssDNA) genome of 4 to 6.3 kilobases (kb) in length [11], and encode at least two major open reading frames (ORFs) that are expressed to produce a non-structural (NS or Rep) and a structural (VP or Capsid) protein.

Parvovirus replication occurs in the nucleus, and can occasionally lead to viral genomes becoming integrated into host chromosomes via non-homologous recombination [12], possibly facilitated by single-stranded breaks created by the nickase function of Rep [12]. Thus, incorporation of parvovirus DNA into the mammalian germline might be expected to occur at a certain frequency, it is also expected that - in the absence of a severe population bottleneck - these newly generated EPV alleles will be rapidly purged from the gene pool via selection and drift. Thus, the presence of numerous independently acquired, fixed EPV insertions in mammalian genomes is unexpected, and suggests that selective pressures may have occasionally favored retention of EPV genes during mammalian evolution. Intriguingly, several EPV loci contain open reading frames (ORFs) capable of expressing complete or almost complete proteins [13, 14], and recent studies have shown that Rep-encoding EPV loci in rodents and elephants exhibit a similar, distinctive pattern of tissue-specific expression (being specifically transcribed in the liver) despite having been acquired in entirely independent germline incorporation events [14].

In this study we use a combination of *in silico* and molecular biological approaches to investigate a guinea pig fusion gene named *enRep-Myo9*, that is partially derived from an EPV, and partially derived from a host gene.

## Results

### Guinea pigs express enRep-Myo9 - a fusion gene partly derived from EPV sequences

A predicted fusion gene containing EPV-derived sequences was identified in the guinea pig (*Cavia porcellus*) via similarity search-based screening of the reference RNA sequence (RefSeq RNA) database [15]. We examined this gene using comparative approaches, revealing it to be comprised of a 3’ truncated parvovirus *rep* gene fused to five exons of a *Myo9*-like gene encoded by guinea pigs (Accession #: XM_013153010.2) (Fig 1). Similarity search-based screening of the rodent whole genome sequence (WGS) assemblies revealed that the viral portions of *enRep-Myo9* are encoded by an EPV locus specific to the cavies (family *Caviidae*,), which we named enRep. Cavies are rodents native to South America, and includes guinea pigs, wild cavies, and maras. The presence of the enRep insertion at an orthologous locus in three guinea pig species, as well as the more distantly related Patagonian mara (*Dolichotis patagonum*) (see Fig S1) demonstrates that it was incorporated into the germline prior to divergence of these species an estimated ~22 million years ago (Mya) [16-18]. Meanwhile, the absence of enRep in more distantly related rodent species, such as the chinchilla (*Chinchilla lanigera*) indicates that germline integration probably occurred less than 35 Mya (Fig 1B). We observed that while most of the enRep locus is highly degraded by mutation (reflecting a long presence in the guinea pig germline), the portions of the Rep reading frame that are incorporated into the coding sequence of the *enRep-Myo9* gene are conserved, consistent with purifying selection.

**Figure 1.**
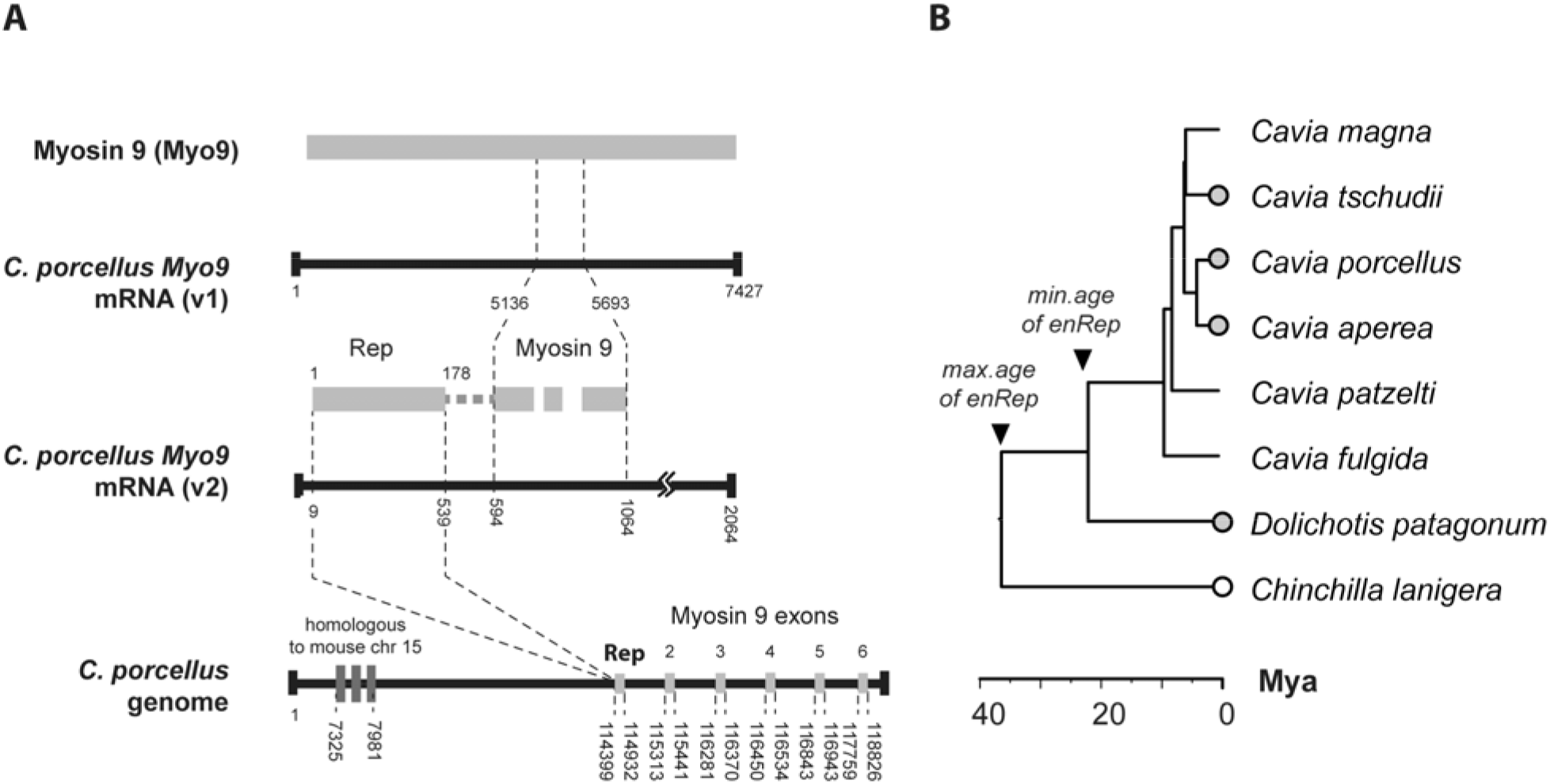
Genomic and phylogenetic characteristics of the *enRep-Myo9* gene. **A.** Schematic diagram showing the structure of the *enRep-Myo9* locus, named as Myosin 9 variant (v2) in the data base, and Myosin 9 (v1) gene product. Black bars indicate nucleic acids (RNA or DNA), grey bars show translated gene products. Numbers show the position of genomic features within each transcript or scaffold in base pairs (bp). Accession numbers of the transcripts shown here are as follows: mRNA version 1(XM_003470459.3); mRNA version 2 (XM_013153010.2); genomic scaffold (AAKN02031205). Abbreviations: Rep=Replicase; chr=chromosome. **B.** A time-scaled phylogenetic tree showing divergence times of six extant species in the genus *Cavia*, and the known distribution of the enRep element in the genus. Circles next to taxa names indicate species examined in this study. Circles with grey fill indicate taxa in which enRep is present. Empty circles indicate taxa in which enRep is absent. All three of these taxa harbor the enRep orthologs in their genomes, indicating that the element was generated more than 22 million years ago (Mya), based on estimated species divergence times.

Investigation of the enRep locus *in silico* revealed that it is located within 0.4Kb of the *Myo9* gene exons that are incorporated into the *enRep-Myo9* gene. We next used polymerase chain reaction (PCR) to amplify; (i) the regions of enRep that are incorporated into the *enRep-Myo9* gene, (ii) their immediate genomic context and; (iii) the full-length gene from genomic DNA (Fig 2A). The resulting amplicons were sequenced, confirming that the locus is structured as in the published WGS assembly. Since computer program-based predictions of genes, transcripts and splicing sites need to be experimentally validated, we generated and sequenced cDNA to confirm that *in vivo* expression and splicing of the *enRep-Myo9* gene occurs as predicted by *in silico* approaches.

**Figure 2.**
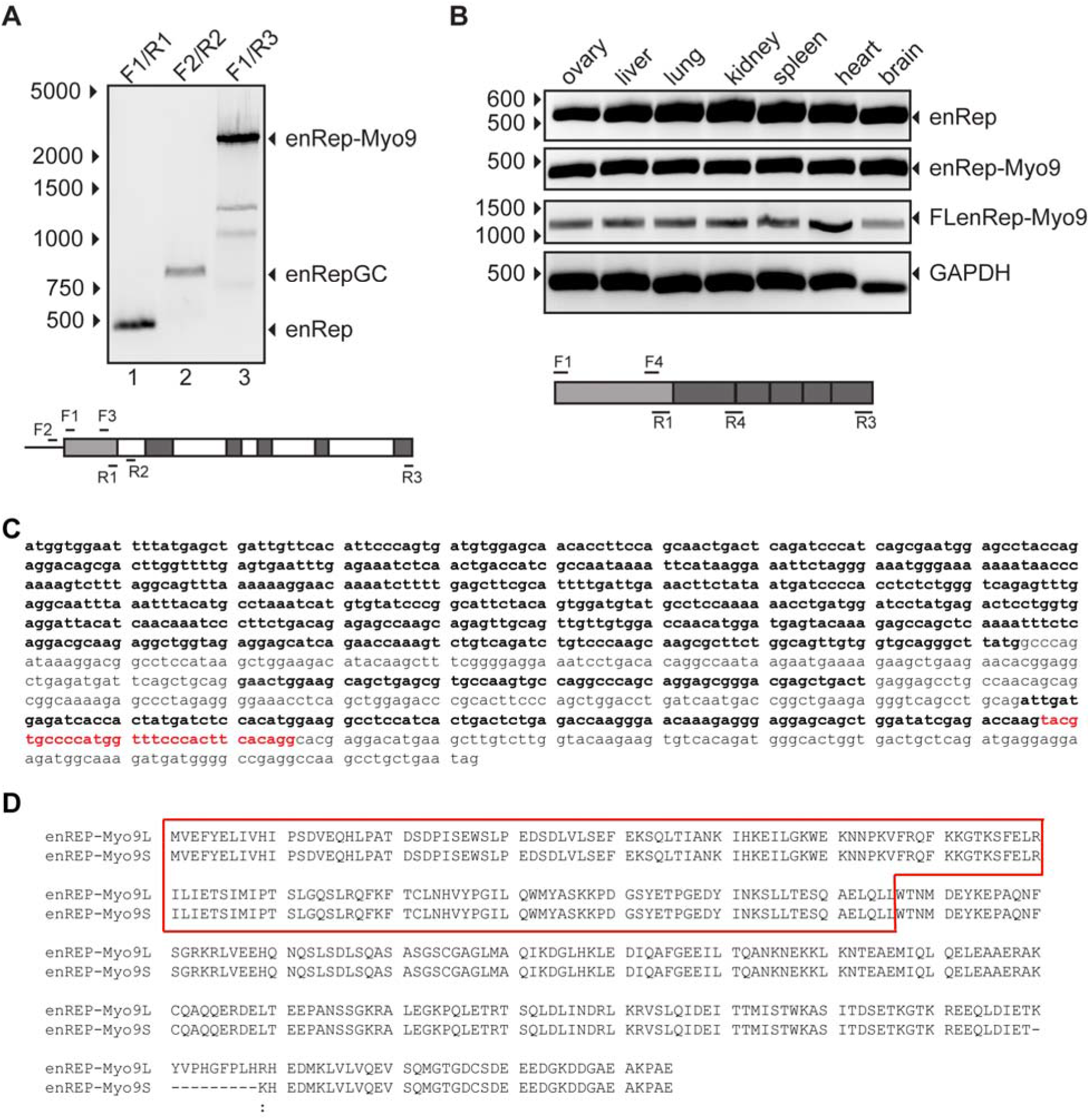
Parvoviral-derived myosin fusion is expressed in guinea pig. **A.** The presence of a parvoviral derived *rep* gene in the genome of guinea pig was verified by PCR from genomic DNA. F1/R1 primers were used to amplify regions of the *rep* gene encoded by an EPV (enRep), F2/R2 primers were used to amplified *rep* in its immediate genomic context (enRepGC). F1/R3 primers were used to amplified *rep* and *Myo9* homologous exons (enRep-Myo9). All amplicons were verified by sequencing. The location of the primers on the gene is shown as follows: *rep* hexon in gray, *Myo9* homologous exons in black, introns in white. **B.** The expression of *enRep-Myo9* transcript was detected in several guinea pig tissues. Total RNA was isolated, cDNA was generated using oligo dT and used for PCR to detect enRep (enRep), *enRep-Myo9* until the exon 2-3 junction (enRep-Myo9), or the full length *enRep-Myo9* coding sequence (FLenRep-Myo9). *Cp*GAPDH mRNA was used as loading control. The location of the primers on the enRep-Myo9 coding sequence is shown below. The DNA ladder migration is indicated on the left-hand side. **C.** The *enRep-Myo9* coding sequence confirmed by sequencing of cDNA is shown. Distinct exons are indicated by bold letters. The sequence missing in the alternatively spliced version is shown in red. **D.** Alignment of the long (enRep-Myo9L) and short (enRep-Myo9S) proteins encoded by the cloned sequences. The enRep domain is framed in red.

### The enRep-Myo9 gene is broadly transcribed in vivo

We investigated *in vivo* transcription of *enRep-Myo9* using PCR. RNA was isolated from several guinea pig (*C.* p*orcellus*) tissues and used to generate cDNA. The absence of genomic DNA in cDNA preparations was confirmed prior to performing assays (data not shown). We first demonstrated that cDNA encoding the enRep section of the *enRep-Myo9* mRNA is present in all tissues analyzed (Fig 2B). Next, we verified that these sequences are expressed as part of a transcript encoding a fusion protein comprised of enRep sequences fused to Myo9 exons as predicted. PCR using a forward primer at the end of enRep, and a reverse primer aligning at the junction of exons 2 and 3 of Myo9 (Fig 2B), generated amplicons of the expected size in all samples analyzed, confirming that *enRep-Myo9* is expressed as a fused transcript.

The presence of the full-length coding sequence was confirmed in all analyzed tissues (Fig 2B): ovary, liver, lung, kidney, spleen, heart and brain by PCR. These last amplicons were sequenced, demonstrating that there are two alternatively spliced forms of *enRep-Myo9*. The isoforms differ at the junction of Myo9 exons 5 and 6, with the shorter isoform lacking a region spanning ten codons of exon 5 (Fig 2D). In some tissue samples (e.g. spleen), both isoforms were found to occur. We failed to obtain the untranslated regions of the mature *enRep-Myo9* transcript, despite several attempts. Thus, we cannot confirm the entire predicted transcript sequence, which includes a long 3’UTR and a very short 5’UTR, is expressed. However, our analysis does confirm that the entire coding sequence is expressed.

### The enRep-Myo9 protein exhibits cytosolic localization in vivo

To verify if the ORFs in the *enRep-Myo9* transcript can be translated into a protein, we cloned and expressed the two isoforms of *enRep-Myo9 in vitro*. enRep-Myo9 long (380 amino acids) and short (370 amino acids) coding sequences were cloned from spleen derived cDNA, into eukaryotic expression vectors to express a N- or C-terminal Flag-tagged protein. Cells were transfected with the 3×Flag-enRep-Myo9L or S, enRep-Myo9-FlagL or S or control plasmids, a Flag-tag protein was observed when either construct was used (Fig 3A) and there were not notable differences in mobility between the isoforms. This shows that it is possible to obtain a protein from the coding sequence present at the *enRep-Myo9* transcripts.

Western blot assays only show a migration difference between the Flag-enRep-Myo9 and enRep-Myo9-Flag proteins, which is due to the three Flag epitopes added when the protein is tagged in the amino terminal. The expected size of both proteins (including the epitopes) was <50kDa, but we observed a migration above the 50 kDa. It is known that parvoviral replicases are regulated by phosphorylation [19], and several programs show putative posttranslational modification in both, short and long enRep-Myo9. Thus, the lower migration observed for enRep-Myo9 might reflect posttranslational modification of the protein.

To further investigate the enRep-Myo9 protein, we asked where in the cell it might be located. Unlike AAV Rep and other EPVs containing intact ORFs encoding complete or nearly complete Rep proteins (e.g. the Odegus4 element in the genome of the degu [*Octodon degus*]) [13], the enREP-Myo9 sequence does not contain nuclear localization signals. We cloned and expressed the Odegus4 element, demonstrating that this transcript can express a protein (Fig 3A). We compared the localization of Odegus4 to enRep-Myo9S expressed from both constructs and include GFP-Myosin9 from mouse in the analysis. As expected, the protein expressed by Odegus4 is mainly located in the nucleus, while GFP-Myosin9 has cytoplasmic localization, both enREP-Myo9 proteins have cytoplasmic localization and no difference is observed between the isoforms (data not shown).

**Figure 3.**
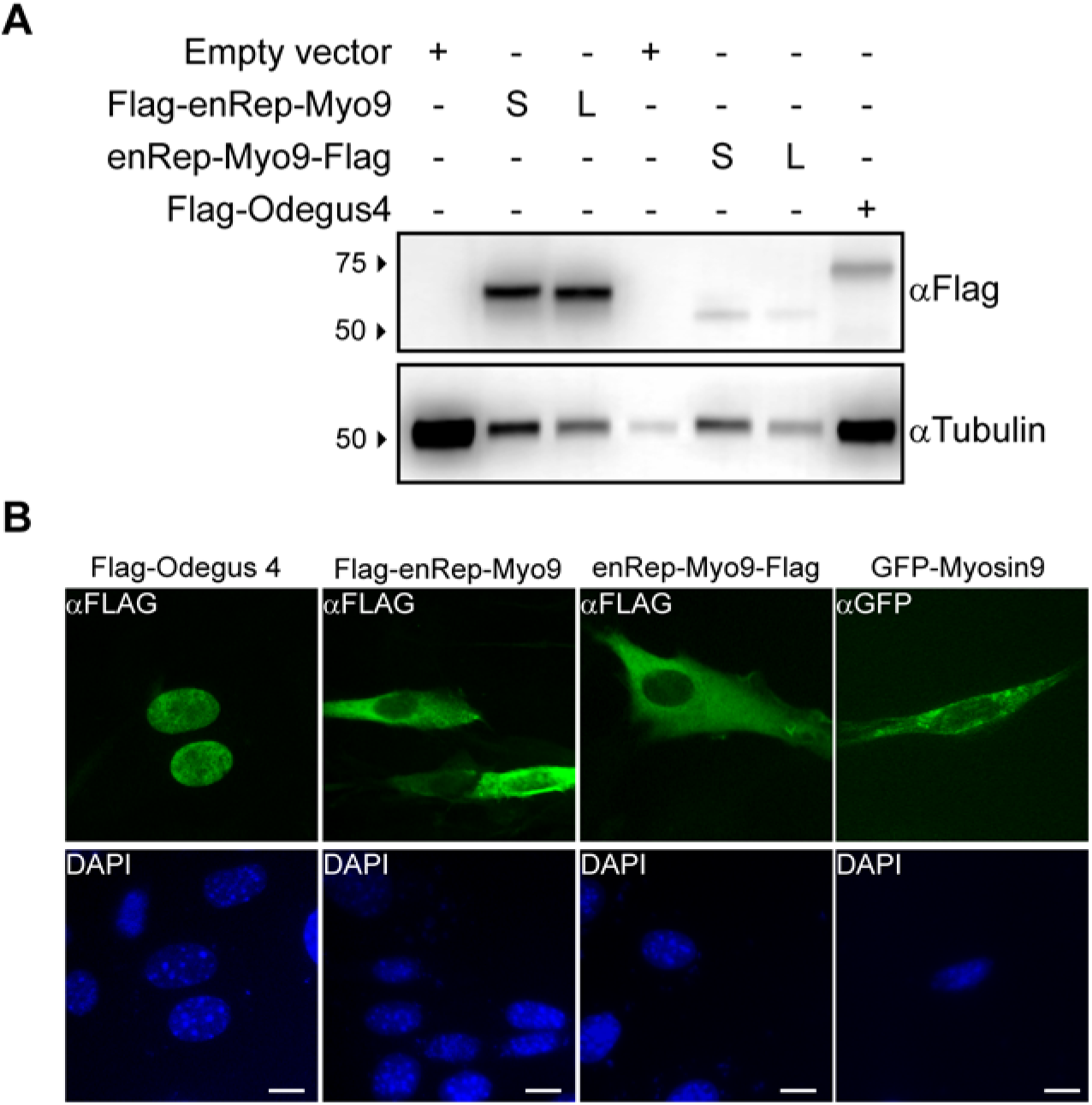
enRep-Myo9 encodes a protein with cytoplasmic localization. The expression of enRep-Myo9 isoforms was analyzed by western blot and immunofluorescence. **A.** NIH3T3 were transfected with plasmids encoding Flag-enRep-Myo9L, Flag-enRep-Myo9S, enRep-Myo9-FlagL, enRep-Myo9-FlagS, Flag-Odegus4 or empty vector. 48h later cells were lysed and western blot assays were performed using anti-Flag or anti-Tubulin antibodies as loading control. The migration of the molecular marker is shown on the left-hand side, the antibody used for each blot is indicated on the right-hand side. **B.** NIH3T3 seeded over coverslips were transfected with plasmids encoding Flag-enRep-Myo9S, enRep-Myo9-FlagS, Flag-Odegus4 or mouse GFP-Myosin9. 24h later cells were fixed and immunofluorescence was performed using anti-Flag or anti-GFP antibodies. DAPI was used to stain the nucleus. Bar size is 10µm.

## Discussion

In this study, we investigated the evolution and molecular properties of a guinea pig fusion gene: *enRep-Myo9*. We show that the viral portions of *enRep-Myo9* derive from an EPV (named enRep) that was incorporated into the genome of caviomorph rodents between 6-35 Mya, while the host portions derive from five exons of a *myosin 9* (*Myo9*)-like gene. Furthermore, while most of this EPV sequence is degraded, the portions included in the *enRep-Myo9* gene are intact in multiple species of guinea pig (genus *Cavia*), consistent with evolution under purifying selection. Using PCR, we confirmed that *enRep-Myo9* is expressed *in vivo* (Fig 2B) and show that in some tissues, two splicing forms are present (Fig 2C).

The enRep-Myo9 protein contains two main domains, the N-terminal Rep_N domain (amino acids 4-176) and the C-terminal myosin tail domain (amino acids 58-325). It was not possible to identify a nuclear localization signal (NLS) for the enRep section of the protein, and consistent with this, we observed cytosolic localization for the enRep-Myo9 protein regardless of whether it was tagged at the N- or C- terminus. We compared the localization of enRep-Myo9 with another EPV-encoded Rep protein: Odegus4, encoded by a dependoparvoviruses-derived EPV in the degu genome. We previously reported that this EPV is specifically transcribed in degu liver [13]. The predicted gene product contains at least two NLS, as it is expected for Rep proteins (which must locate to the nucleus to allow replication of parvoviral DNA). We demonstrate that the Odegus4 protein indeed exhibits nuclear localization. Thus, if both EPV-derived (and partially EPV-derived) gene products have functional roles, they are likely to be entirely different. We hypothesize that during the adaptation of enRep to its ancestral host the NLS was lost as part of the adaptation process, allowing it to be co-opted for a still unknow cellular role at the cytoplasm. The observation that *enRep-Myo9* is expressed in all analyzed tissues (ovary, liver, lung, kidney, spleen, heart and brain) strongly support the idea that this fusion has an important functional role in guinea pig.

Several EVEs encoding intact open reading frames have been identified in mammalian genomes, and some of these have been studied experimentally, were researchers have cloned the coding sequences and expresses the encoded proteins in heterologous systems to show different functions. Many EVE have been co-opted to perform antiviral activities, the proteins encoded by EVEs can block different steps of viral infection or they can directly regulate the immune system (reviewed in [3, 7]). Also, EVE derived non-coding RNAs may act in DNA regulation as well as interferent RNAs (reviewed in [3, 7]). Most examples come from endogenous retroviruses (ERVs) and only little information exist for co-option of non-retroviral EVE, been endogenous borna-like virus (EBLs) the only example so far. Bornaviruses are enveloped, nonsegmented, negative-strand RNA viruses, that replicate in the nucleus, where the mRNA of their different genes has integrated in different animals, including humans [1, 20]. The main EVEs derived from bornavirus are the endogenous borna-like nucleoprotein (EBLN), the EBLN encoded by the thirteen-lined ground squirrel (*Ictidomys tridecemlineatus*) was shown to colocalize in the nucleus with viral factories and inhibit *in vitro* replication of borna disease virus (BDV) [6], while the protein encoded by human EBLN-1 showed cytoplasmic localization and was unable to inhibit BDV replication [6]. In humans EBLN-1 is expressed in testis and brain [21], but regardless of the fact that has an open reading frame, it is propose to function in gene regulation as a non-coding RNA (ncRNA), rather than as a protein [7, 21]. We do not have yet antibodies that can confirm or discard that *enRep-Myo9* is expressed as protein, but our data do not suggest that enRep-Myo9 works as a regulatory RNA: the fact that we were able to amplify the coding sequence from cDNA generated using oligo dT from several tissues is a strong evidence that this is an expressed mRNA. Also, both splicing forms have ORFs.

The selective forces underlying the fixation of EPVs (and EVEs derived from other kinds of virus) remain poorly understood. Nonetheless, it is now clear that sequences acquired from viruses can be ‘exapted’ by animal genomes in a variety of ways [1, 3, 7, 20]. Our findings indicate that in guinea pigs, a novel gene may have arisen through fusion of EPV and host genes. If indeed this is the case, it would represent the first example of a co-opted/exapted EPV. We demonstrate that the *enRep-Myo9* possesses molecular characteristics that are at least consistent with a functional role as a coding gene, being conserved, broadly transcribed, and capable of expressing protein. These findings provide indications that acquisition of genes from parvoviruses has shaped the evolution of mammals and establishes a platform for future experiments designed to determine whether *enRep-Myo9* plays a biologically relevant role in guinea pig physiology.

## Materials and Methods

### Genome screening in silico

Whole genome sequence (WGS) data were obtained from the National Center for Biotechnology Information (NCBI) genomes resource. WGS data of 60 rodent species (Table S1) were screened for parvoviral-like sequences using the database-integrated genome-screening (DIGS) tool [22]. Parvovirus protein sequences were used as “probes” in similarity search-based screens of WGS data. Sequences that produced statistically significant matches to probes were extracted and classified by BLAST-based comparison to a set of virus reference genomes.

### Nucleic acid extractions and PCR

Two female individuals of 300g were euthanized by 1-cloro-2,2,2-trifluoroetil difluorometil eter overdose according to the protocol approved by the Bioethical Committee of Facultad de Ciencias de la Vida, Universidad Andres Bello (Acta 010/2016). Genomic DNA was obtained from liver tissue, total RNA was extracted from ovary, lungs, liver, kidney, spleen, heart and brain using RNaesy mini kit (Qiagen). cDNA was synthesized using 500 ng of RNA, oligo dT and SuperScript III First-Strand Synthesis System (Thermo Fisher).

To verify the presence of enRep in the genomic DNA, PCR was performed using the primers F1 5’atggtggaattttatgagctg3’ and R1 5’cataagccctgcaccacaactg3’ with 75ng of gDNA, GoTaq Flexi G2 (Promega), 2.0 mM MgCl2 with a program of 95°C, 3 min; 35×[95°C, 30 s; 53.3°C, 30 s; 72°C, 35 s] and a final extension at 72°C, 5 min. To confirm that enRep was in the correct genomic context we use primers aligning 165bp upstream (F2 5’aacctgagttgtcattcagg3’) and 55bp downstream (R2 5’tgatgcccttctgtatgagg3’) of enRep with the same conditions as before except 35×[95°C, 30 s; 53.3°C, 30 s; 72°C, 45 s]. To verify the full *enRep-Myo9* exon-introns in the genome, we preformed PCR using PFUultra II (Agilent), the primers F1 and R3 5’gaggtgggcagtcatccatc3’, that align in the last *Myo9* exon, a program of 92°C, 2 min, 35× [92°C, 20 s; 60°C, 30 s; 72°C, 1 m 50 s] and a final extension at 72°C, 3 min. All the amplicons were cloned in pGEMTeasy (Promega) and sequenced.

To discard genomic DNA contamination in the cDNA preparations, the primers F3 5’tggatcctatgagactcctggt3’ and R2 were used for a PCR reaction with 1µl of cDNA, GoTaq Flexi G2 (Promega), 2.0 mM de MgCl2, and a program of 95°C, 3 min; 35× [95°C, 30 s; 57°C, 30 s; 72°C, 45 s] with a final extension of 72°C, 5 min. To amplify enRep from cDNA the same primers and condition for enRep in genomic DNA were used, except that 40 PCR cycles were performed. To verify the presence of the *enREP-Myo9* transcript a forward primer aligning at the 3’of enRep (F4 5’atggtggaattttatgaggtg3’) and a reverse primer aligning at the exon-exon junction of *enRep-Myo9* exons 2 and 3 (R4 5’cataagccctgcagctgaatc3’) were used under the same conditions as before except 40×[95°C, 30 s; 58°C, 30 s;72°C, 20 s]. For GAPDH (Accession # NM_001172951) the primers Frw 5’gaatcacgagaagtacgaca3’ and Rev 5’gtatttggccggtttctcc3’were used under the same conditions except 40×[95°C, 30 s; 57.3°C, 30 s;72°C, 25 s]. The full length *enRep-Myo9* coding sequence was amplified using PFUII Ultra from the cDNA, using the primers Frw 5’atatctcgagctattcagcaggcttggcctcgg3’ and Rev 5’atatgaattcaatggtggaattttatgagctg3’, 1µl of cDNA, and a program of 92°C, 2 min, 40× [92°C, 20 s; 60°C, 30 s; 72°C, 50 s] and a final extension at 72°C, 3 min. The amplicons were cloned in pGEMTeasy and sequenced.

### Cloning and plasmids

pGEMTeasy-enRep-Myo9 containing the amplicons from spleen were digested with *EcoR*I and *Xho*I and cloned into pcDNA3×Flag to generate pcDNA3×Flag-enREP-Myo9L and S, that encodes for 3×FLAG-enRep-Myo9 isoforms. The coding sequences were also amplified from pGEMTeasy-enRep-Myo9 using the primers Frw 5’atatgaattcgccaccatggtggaattttatg3’ and Rev 5’atatctcgagttcagcaggcttggcctcgg3’ using the same PCR conditions as above. The PCR products were digested with *EcoR*I and *Xho*I and cloned into pCMVtag4C to generate pCMVTag4C-enRep-Myo9 L and S, that encodes for enRep-Myo9-Flag isoforms. Odegus4 coding sequence was amplified using the primers Frw 5’atggaattcatggtgcagttttatgagc3’ and Rev 5’atctcgagctagagggcgcactttttcc3’from degu genomic DNA. The PCR product was digested with *BamH*I and *Xho*I and cloned into pcDNA3×Flag to generate pcDNA3×Flag-Odegus4, that encodes for 3×Flag-Odegus4. The plasmid MyosinIIA-GFP [23] that encodes for mouse GFP-Myosin9 was a gift from Matthew Krummel (Addgene plasmid #38297).

### Western blot and immunofluorescence

NIH3T3 cells were seeded in 12 well plates and transfected with 500 ng of pcDNA3×Flag-enRep-Myo9L, pcDNA3×Flag-enRep-Myo9S, pCMVtag4C-enRep-Myo9L, pCMVtag4C-enRep-Myo9S, pcDNA3×Flag-Odegus4 or empty vector. 48 h later the cells were lysed in Reporter lysis buffer (Promega). Samples were then boiled in 5× sodium dodecyl sulphate (SDS) loading buffer, and the proteins were resolved by 10% acrylamide SDS-PAGE. After transfer to nitrocellulose membranes, the blots were probed with mouse anti-Flag (Clone M2, Sigma) or mouse anti-α tubulin (Clone DM1A, Sigma). Secondary antibodies conjugated to HRP and the ECL reagents were used for developing.

For immunofluorescence 2,5×10^4^ NIH3T3 seeded in 12mm coverslip, were transfected with 500 ng of pcDNA3×Flag-enRep-Myo9S, pCMVtag4C-enRep-Myo9S, pcDNA3×Flag-Odegus4 or pEGFP-Myosin9. 48 hour later were fixed with 3.7% formaldehyde in phosphate buffered saline. After permeabilization the covers were incubated with mouse anti-Flag M2 1:100 or mouse anti-GFP B2 (Santa Cruz Biotechnology) 1:100. Secondary antibodies anti-mouse AlexaFluor-488 1:500 was used. Samples were mounted in prolong diamond fade with 4′,6-diamidino-2-phenylindole (DAPI) and fluorescent images were acquired at ×40 magnification with DFC3000G Leica charge-couple-device (CCD) camera mounted on Leica DMIL microscope running Leica LASX software. Digital images were processed with Photoshop CS3 (Adobe).

## Acknowledgments

This work was supported by Fondo Nacional de Desarrollo Científico y Tecnológico Grant FONDECYT1180705 to GA. IVH was supported by AAP2018-1 from Universidad Andres Bello. RJG was funded by the Medical Research Council of the United Kingdom (MC_UU_12014/12).

## References

1. Katzourakis, A. and R.J. Gifford, Endogenous viral elements in animal genomes. PLoS Genet, 2010. 6(11): p. e1001191.

2. Feschotte, C. and C. Gilbert, Endogenous viruses: insights into viral evolution and impact on host biology. Nat Rev Genet, 2012. 13(4): p. 283–96.

3. Frank, J.A. and C. Feschotte, Co-option of endogenous viral sequences for host cell function. Curr Opin Virol, 2017. 25: p. 81–89.

4. Gautam, P., T. Yu, and Y.H. Loh, Regulation of ERVs in pluripotent stem cells and reprogramming. Curr Opin Genet Dev, 2017. 46: p. 194–201.

5. Horie, M. and K. Tomonaga, Paleovirology of bornaviruses: What can be learned from molecular fossils of bornaviruses. Virus Res, 2018.

6. Fujino, K., et al., Inhibition of Borna disease virus replication by an endogenous bornavirus-like element in the ground squirrel genome. Proc Natl Acad Sci U S A, 2014. 111(36): p. 13175–80.

7. Horie, M., The biological significance of bornavirus-derived genes in mammals. Curr Opin Virol, 2017. 25: p. 1–6.

8. Kapoor, A., P. Simmonds, and W.I. Lipkin, Discovery and characterization of mammalian endogenous parvoviruses. J Virol, 2010. 84(24): p. 12628–35.

9. Belyi, V.A., A.J. Levine, and A.M. Skalka, Sequences from ancestral single-stranded DNA viruses in vertebrate genomes: the parvoviridae and circoviridae are more than 40 to 50 million years old. J Virol, 2010. 84(23): p. 12458–62.

10. Liu, H., et al., Widespread endogenization of densoviruses and parvoviruses in animal and human genomes. J Virol, 2011. 85(19): p. 9863–76.

11. Cotmore, S.F., et al., The family Parvoviridae. Archives of virology, 2014. 159(5): p. 1239–1247.

12. Berns, K.I., Parvovirus replication. Microbiological reviews, 1990. 54(3): p. 316–329.

13. Arriagada, G. and R.J. Gifford, Parvovirus-derived endogenous viral elements in two South American rodent genomes. J Virol, 2014. 88(20): p. 12158–62.

14. Kobayashi, Y., et al., An endogenous adeno-associated virus element in elephants. Virus Res, 2019. 262: p. 10–14.

15. O’Leary, N.A., et al., Reference sequence (RefSeq) database at NCBI: current status, taxonomic expansion, and functional annotation. Nucleic Acids Res, 2016. 44(D1): p. D733–45.

16. Dunnum, J.L. and J. Salazar-Bravo, Molecular systematics, taxonomy and biogeography of the genus Cavia (Rodentia: Caviidae). Journal of Zoological Systematics and Evolutionary Research, 2010. 48(4): p. 376–388.

17. Opazo, J.C., A molecular timescale for caviomorph rodents (Mammalia, Hystricognathi). Mol Phylogenet Evol, 2005. 37(3): p. 932–7.

18. Fabre, P.H., et al., A glimpse on the pattern of rodent diversification: a phylogenetic approach. BMC Evol Biol, 2012. 12: p. 88.

19. Nuesch, J.P., et al., Regulation of minute virus of mice NS1 replicative functions by atypical PKClambda in vivo. J Virol, 2003. 77(1): p. 433–42.

20. Horie, M. and K. Tomonaga, Non-retroviral fossils in vertebrate genomes. Viruses, 2011. 3(10): p. 1836–48.

21. Sofuku, K., et al., Transcription Profiling Demonstrates Epigenetic Control of Non-retroviral RNA Virus-Derived Elements in the Human Genome. Cell Rep, 2015. 12(10): p. 1548–54.

22. Zhu, H., et al., Database-integrated genome screening (DIGS): exploring genomes heuristically using sequence similarity search tools and a relational database. bioRxiv, 2018: p. 246835.

23. Jacobelli, J., et al., Myosin-IIA and ICAM-1 regulate the interchange between two distinct modes of T cell migration. J Immunol, 2009. 182(4): p. 2041–50.

